# SPPIDER-seq: Sequence-based partner-aware predictor of protein-protein interaction sites

**DOI:** 10.64898/2026.04.28.721359

**Authors:** Aleksey Porollo, Om Jadhav, Aaron Alvarez, Jichao Chen

## Abstract

**Motivation:** Sequence-based protein–protein interaction (PPI) site predictors typically analyze proteins in isolation, neglecting partner-specific context that is critical for interface specificity, particularly in transient and disordered interactions.

**Results:** We introduce SPPIDER-seq, a partner-aware PPI site prediction framework that combines pretrained ESM-2 embeddings with a cross-attention architecture to enable residue-level conditioning on interacting partners. We curated non-redundant protein–peptide interaction datasets from BioLiP and used them to train and benchmark two complementary models: a receptor-centric model optimized for structured interfaces and a peptide-centric model tailored to disordered, motif-driven binding. On blind benchmarks, SPPIDER-seq achieved AUROC values up to 0.797 and MCC values up to 0.269, outperforming AlphaFold3 on peptide-mediated and disordered interfaces while remaining complementary on globular complexes. Application to 341 TP53 interaction partners revealed coherent, partner-specific interface patterns across both structured and intrinsically disordered regions.

**Availability and Implementation:** SPPIDER-seq models, datasets, and the Python code are freely available on the web at: https://github.com/aporollo-lab/SPPIDER-seq and archived on Zenodo at DOI: 10.5281/zenodo.19835990, corresponding to GitHub release v2.0-manuscript.

**Contact:** Dr. Aleksey Porollo –

Dr. Jichao Chen –

**Supplementary Information:** Available online with the manuscript.

## 1. Introduction

Protein–protein interactions are integral to virtually every biological process, including signal transduction, immune regulation, cell cycle progression, and metabolic control. Accurately identifying PPI sites at the residue or atomic level is critical for elucidating the molecular basis of cellular mechanisms, interpreting the impact of disease-associated mutations, and guiding the rational design of targeted therapeutics and biologics.

Experimental techniques have been instrumental in mapping protein–protein interaction (PPI) interfaces. Alanine scanning mutagenesis remains a classical method for identifying energetic hot spots (Bogan & Thorn, 1998; Moreira et al., 2007), while X-ray crystallography has provided atomic-resolution views of protein complexes, enabling precise localization of key interface residues (Chakrabarti & Janin, 2002; Jones & Thornton, 1996). Nuclear magnetic resonance (NMR) spectroscopy complements these methods by probing transient interactions and conformational dynamics, particularly in disordered or flexible regions (Sattler et al., 1999; Vaynberg & Qin, 2006). More recently, cryo-electron microscopy (Cryo-EM) has emerged as a powerful tool for resolving large, dynamic, and membrane-associated assemblies without crystallization requirements (Callaway, 2015; Nogales & Scheres, 2015). Despite their contributions, these methods face limitations in cost, throughput, and applicability to unstable or flexible proteins. These constraints highlight the need for robust, scalable computational approaches capable of efficiently predicting PPI sites using sequence-derived features, structural context, and evolutionary signals to complement and extend experimental insights.

Over the past two decades, computational approaches for predicting PPI sites have advanced considerably. Early efforts primarily relied on manually engineered features extracted from protein sequence and structure, such as solvent accessibility, residue depth, protrusion index, evolutionary conservation, electrostatic potential, and hydrophobicity profiles. These features were typically input into traditional machine learning classifiers, including support vector machines (SVMs), random forests, and naïve Bayes models, which formed the foundational architecture of early predictive frameworks aimed at estimating residue-level interaction propensities. These models often required extensive feature selection and domain-specific expertise but enabled meaningful predictions even in the absence of high-resolution structures. The reader may refer to comprehensive reviews of representative methods and their applications in protein interface prediction in (Kiouri et al., 2025; Xue et al., 2015; Zhou & Qin, 2007).

The emergence of deep learning significantly transformed PPI site prediction by enabling automatic feature extraction from raw input data, such as amino acid sequences, residue graphs, or voxelized structural representations. Architectures such as convolutional neural networks (CNNs) or graph neural networks (GNNs) facilitated the modeling of intricate spatial and sequential dependencies that traditional machine learning approaches struggled to capture. Importantly, deep learning approaches proved valuable for identifying interface residues even in proteins with limited structural annotations by leveraging local and global contextual cues. For detailed overviews of deep learning-based interface prediction methods prior to the advent of large-scale transformer models, the reader is referred to the reviews in (Kiouri et al., 2025; Tang et al., 2023). However, these models often required large volumes of high-quality annotated data for training and were sensitive to noise in predicted structures or incomplete sequence features. Moreover, generalization across diverse protein folds and interaction types remained challenging, particularly in the absence of partner-specific information. Despite these limitations, early deep learning methods laid a strong foundation for the current wave of end-to-end representation learning techniques based on contextual embeddings, which now dominate the field.

A transformative leap in protein interface prediction has come with the adoption of transformer-based protein language models (PLMs), architectures originally developed for natural language processing (NLP), to decipher the underlying “grammar” of protein sequences. Models such as ProtT5 (Elnaggar et al., 2022) and ESM-2 (Lin et al., 2023), trained on hundreds of millions of sequences, have demonstrated an unprecedented ability to learn rich representations that implicitly encode evolutionary, structural, and functional information. These embeddings have been successfully repurposed for diverse downstream tasks, including protein structure prediction (Gomez-Uribe et al., 2025; Madani et al., 2023; Ruffolo & Madani, 2024; Z. Zhang et al., 2024) or mutation effect analysis (Marquet et al., 2022; Marquet et al., 2024; Sun & Shen, 2025). Notably, recent studies have applied ProtT5 and ESM-2 embeddings to residue-level classifiers for predicting interface residues, achieving state-of-the-art performance while bypassing the need for hand-crafted features or explicit structural inputs (Littmann et al., 2021; Sargsyan & Lim, 2024). This convergence of language modeling and structure prediction signals a paradigm shift toward highly accurate, context-sensitive prediction of PPI sites at proteome scale.

Despite significant progress in PPI site prediction, several critical challenges remain unresolved. One major limitation of many existing computational approaches is their lack of partner-awareness – they typically treat the target protein in isolation, neglecting the contextual influence of its binding partner. This omission restricts the ability of models to accurately delineate partner-specific interfaces, particularly in transient PPIs where interaction sites are often small, dynamic, and condition-dependent. The problem is further exacerbated in intrinsically disordered regions (IDRs), which lack stable tertiary structures and exhibit lower sequence conservation. These regions frequently mediate transient interactions via flexible binding motifs, making them poorly suited to structure-based or conservation-driven predictors. Furthermore, existing methods, both experimental and computational, often overlook small binding sites, which are crucial in regulatory signaling and cellular dynamics. Notably, the recently introduced AlphaFold3 predicts multimeric macromolecule structures with atomic resolution, effectively delineating interaction interfaces without explicit interface annotation, but the accurate modeling of heterogeneous conformations and partner-dependent interfaces within disordered regions remains challenging (Abramson et al., 2024).

Addressing these issues, we present a novel partner-aware PPI site prediction framework that leverages a cross-attention mechanism layered atop the ESM-2 pretrained protein language model. By enabling dynamic information exchange between the target protein and its interacting partner, our model captures inter-protein dependencies and adapts to the context of the specific interaction, thus offering improved precision in interface localization. This architecture represents a step forward in building interpretable, high-resolution predictive tools capable of handling complex scenarios such as IDR-mediated interactions and context-dependent interface plasticity.

## 2. Methods

### 2.1. Model Architecture and Training Procedure

To predict PPI sites, a partner-aware deep learning model was developed, incorporating a cross-attention mechanism applied to embeddings from the pretrained ESM-2 protein language model (Lin et al., 2023). Specifically, embeddings were generated using the esm2_t33_650M_UR50D checkpoint, which produces 1280-dimensional per-residue representations. For each interacting protein sequence *S*_*A*_ and *S*_*B*_, the corresponding embedding matrices were defined as:

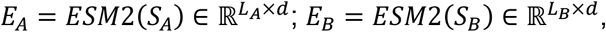

where *L*_*A*_ and *L*_*B*_ denote sequence lengths and *d*=1280 is the embedding dimension.

To explicitly model inter-sequence dependencies, a unilateral multi-head cross-attention layer with 16 attention heads was applied. Attention weights were computed as:

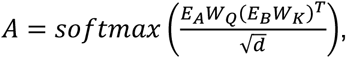

where *W*_*Q*_ and *W*_*K*_ are learnable projection matrices. The resulting matrix 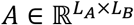 encodes how each residue in protein *S*_*A*_ (query) attends to residues in protein *S*_*B*_(context).

The cross-attended contextual embeddings of a query were then updated as:

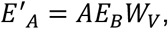

where *W*_*V*_ is a learnable value projection. This attention module captures inter-protein dependencies by learning context-aware interactions at the residue level.

The cross-attended embeddings were passed through a layer normalization step and a two-layer feedforward network, consisting of a linear projection, ReLU activation, dropout, and a final sigmoid output layer. Per-residue interaction probabilities for a query protein were obtained as:

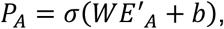

where *W*and *b* are learnable parameters and *σ* is the sigmoid function.

Training on full-length sequences introduces additional challenges, particularly for very large proteins. The underlying protein language model used here (ESM-2) has an effective input length limit of 1,022 amino acids (1024 tokens including BOS and EOS). To accommodate long proteins, we implemented a chunk-wise training strategy, in which sequences were divided into overlapping 1,022-residue segments with a stride of 512. Each segment of the query protein was cross-attended against similarly chunked partner embeddings, and predictions across overlapping windows were aggregated via max pooling. This ensured that predictions remained consistent across sequence boundaries while preserving contextual dependencies.

Training was performed using binary cross-entropy loss, with residue-level labels denoting interface (positive) and non-interface (negative) residues. Performance was monitored on a held-out validation subset, and early stopping was applied if validation loss did not improve for 10 consecutive epochs. Training was limited to a maximum of 50 epochs, and the model parameters corresponding to the lowest validation loss were retained for final evaluation on a blind test set.

To account for the distinct patterns of interaction site distribution observed between receptor proteins and short peptide ligands, two separate models were trained: one *receptor-centric* and the other *peptide-centric*. This distinction reflects structural and functional asymmetries in their interaction interfaces and is further examined in Supplementary Materials SD1.

### 2.2. Datasets

Due to the limited availability of experimentally validated PPI data involving intrinsically disordered regions, we used receptor–peptide complexes from the BioLiP database (C. Zhang et al., 2024) as a large source of experimentally resolved protein–peptide interaction interfaces, which include both structured and disorder-mediated binding modes. BioLiP is a curated resource that extracts biologically relevant ligand–protein interactions from the Protein Data Bank (PDB) (Berman et al., 2000; Burley et al., 2025) and includes peptide ligands of ≤30 residues, making it particularly suitable for modeling short, flexible binding motifs.

Stringent filtering criteria were applied to ensure biological relevance and sequence integrity. Complexes were excluded if they (i) lacked a UniProt reference, (ii) mapped to multiple UniProt identifiers, or (iii) showed <90% sequence identity to the corresponding UniProt entry, thereby removing synthetic, chimeric, or misannotated sequences. Antibody–epitope complexes were also excluded, as epitopes are immune recognition targets rather than physiologically evolved binding partners, and their inclusion could bias the dataset toward non-representative structural motifs.

Two non-redundant overall datasets were then constructed: one receptor-centric (dubbed RC-Overall) and the other peptide-centric (PC-Overall). In each dataset, the focal proteins (whose interaction sites were to be predicted) were clustered such that no two receptors or ligands, respectively, shared more than 25% pairwise sequence identity based on the full-length UniProt parent proteins corresponding to the experimentally resolved receptor- and peptide-containing chains. No explicit reciprocal alignment coverage constraints were applied. As a result, cluster assignment was more inclusive with respect to locally similar sequence regions, leading to a more stringent removal of potentially redundant sequences than would be obtained under additional query/target coverage requirements. The resulting PPI pairs were partitioned into training (60%), validation (10%), and blind test (30%) subsets (**Table 1**). If a focal protein participated in multiple PPI pairs with different partners, all corresponding pairs were assigned to the same subset, thereby preventing data leakage across splits.

**Table 1.**
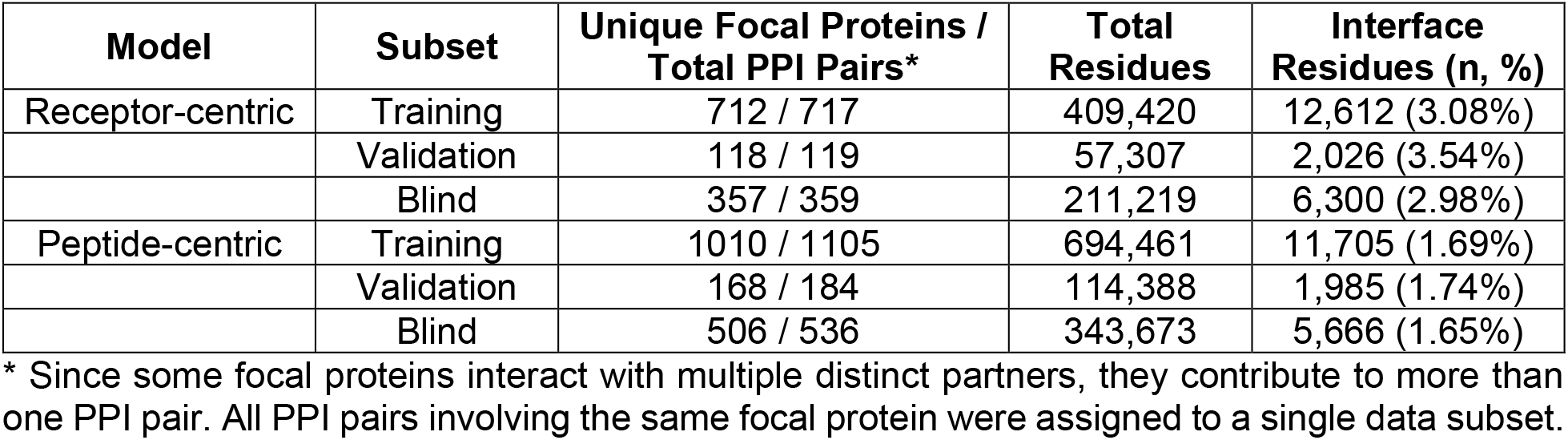
Dataset composition and class distribution in focal partners across training, validation, and blind (60:10:30 split) subsets for receptor- and peptide-centric models.

| Model | Subset | Unique Focal Proteins / Total PPI Pairs* | Total Residues | Interface Residues (n, %) |
| --- | --- | --- | --- | --- |
| Receptor-centric | Training | 712 / 717 | 409,420 | 12,612 (3.08%) |
|  | Validation | 118 / 119 | 57,307 | 2,026 (3.54%) |
|  | Blind | 357 / 359 | 211,219 | 6,300 (2.98%) |
| Peptide-centric | Training | 1010 / 1105 | 694,461 | 11,705 (1.69%) |
|  | Validation | 168 / 184 | 114,388 | 1,985 (1.74%) |
|  | Blind | 506 / 536 | 343,673 | 5,666 (1.65%) |

\* Since some focal proteins interact with multiple distinct partners, they contribute to more than one PPI pair. All PPI pairs involving the same focal protein were assigned to a single data subset.

To assess robustness with respect to data partitioning, in addition to the single train/validation/blind split, we also conducted a 5-fold grouped cross-validation. Grouping was performed at the focal UniProt level (receptor-centric or peptide-centric, depending on the model), ensuring that no focal protein or peptide sequence appeared across folds. In each iteration, one fold (20% of the data) was held out as a blind test set, while the remaining four folds (80%) were further partitioned into training (70%) and validation (10%) subsets, yielding an overall 70:10:20 ratio for training, validation, and blind sets in each fold.

Interface residue labels were assigned using experimental structural data and a previously established method (Porollo & Meller, 2007). A residue was annotated as an interface site if its relative solvent accessibility (RSA) changed by >4% upon separation of the complex into individual chains. This RSA-based criterion ensured consistent annotation across all structures. Identified sites were then mapped onto full-length UniProt sequences, and only complete sequences were retained for model training, validation, and downstream benchmarking.

To assess the fraction of disordered ligands in the peptide-centric blind subset, we retrieved full-length 3D models from the AlphaFold2 (AF2) database (Fleming et al., 2025; Jumper et al., 2021) and analyzed their secondary structure profiles. Peptide ligands containing ≥75% unstructured coil (extended conformation) residues were classified as disordered. AF2 models of single chain proteins were used instead of BioLiP experimental structures to account for potential induced-fit effects, whereby intrinsically unstructured peptides adopt ordered conformations upon binding (Csermely et al., 2010; Goh et al., 2004; Trellet et al., 2013). Of 536 pairs analyzed, 212 contained PPI interfaces localized within disordered regions in peptide ligands.

For benchmarking against AlphaFold3 (AF3) (Abramson et al., 2024), we noted that AF3 failed to process a small subset of blind test pairs due to either the 5000-token input limitation of the AF3 web server or unspecified job failures. Specifically, AF3 successfully processed 343 of 359 receptor-centric pairs and 527 of 536 peptide-centric pairs. Among the 212 peptide-centric disordered-peptide cases, 209 were successfully processed. For internal evaluations, we therefore used the complete blind test sets named as follows: RC-BS359 (receptor-centric), PC-BS536 (peptide-centric), and PC-DP-BS212 (disordered peptides in the peptide-centric blind set). For direct AF3 comparisons, only successfully processed subsets were used, named RC-AF3-BS343, PC-AF3-BS527, and PC-AF3-DP-BS209, respectively.

In addition to benchmarking against AF3, we further compared our model performance with a sequence-based framework for intrinsic disorder IUPred2A and its companion method ANCHOR2 (Meszaros et al., 2018) for binding region prediction using the same receptor- and peptide-centric blind datasets. Notably, a recent review by Basu *et al*. (Basu et al., 2023) ranks IUPred2A/ANCHOR2 among the top-performing methods, citing results from the Critical Assessment of Intrinsic Disorder Prediction (CAID) experiment (Necci et al., 2021). This enabled comparison with a widely used baseline specifically designed for intrinsically disordered region (IDR)-mediated interactions.

IUPred2A was used to estimate intrinsic disorder under multiple configurations: the “long disorder” and “short disorder” modes for the peptide-centric dataset, and the “structured domains” option for the receptor-centric dataset. The resulting disorder profiles were combined with ANCHOR2 predictions using its default context-dependent mode to identify putative binding regions within disordered segments. To specifically assess performance in disorder-enriched settings, the peptide-centric blind set was further restricted to ligands with ≥75% predicted disorder according to IUPred2A. Using the long and short disorder modes yielded two partially overlapping subsets of 181 proteins each, hereafter referred to as PC-IA-DPld-BS181 and PC-IA-DPsd-BS181, respectively.

Further detailed analysis of the datasets and respective PPI interfaces can be found in Supplementary Materials SD1.

### 2.3. Evaluation Metrics

The performance of the PPI site prediction models was assessed using a suite of binary classification metrics computed at the per-residue level. Unless otherwise specified, a probability cutoff of 0.5 was applied to convert predicted probabilities into binary labels, with residues classified as either interface (positive) or non-interface (negative). The following metrics were used: Accuracy, Recall (Sensitivity), Precision (Positive Predictive Value), F1-score, Matthews Correlation Coefficient (MCC), Area Under the Receiver Operating Characteristic Curve (AUROC), and Area Under the Precision–Recall Curve (PRAUC). Formal definitions of these metrics are provided in Supplementary Materials SD2. Performance variability across data partitions was quantified by reporting the mean and standard deviation of evaluation metrics (Accuracy, Precision, Recall, F1-score, and MCC) across the five folds, providing an estimate of model stability and generalizability under different non-overlapping data splits.

## 3. Implementation

### 3.1. Cross-Attention Model: Concept, Capacity, and Learned Statistics

Proteins frequently interact with different partners using distinct and often non-overlapping regions of their sequence or 3D structure. To accommodate this partner specificity, we developed a cross-attention architecture in which the prediction for a given residue explicitly depends on the identity and sequence context of its interaction partner. As illustrated in **Figure 1A**, residue-level embeddings of Protein A (query) and Protein B (partner) are provided as input. A cross-attention module enables each residue in Protein A to attend to all residues in Protein B, thereby producing context-aware embeddings that encode inter-protein dependency patterns. These attention-refined representations are then processed by a multilayer perceptron (MLP) that outputs residue-level interaction probabilities.

**Figure 1.**
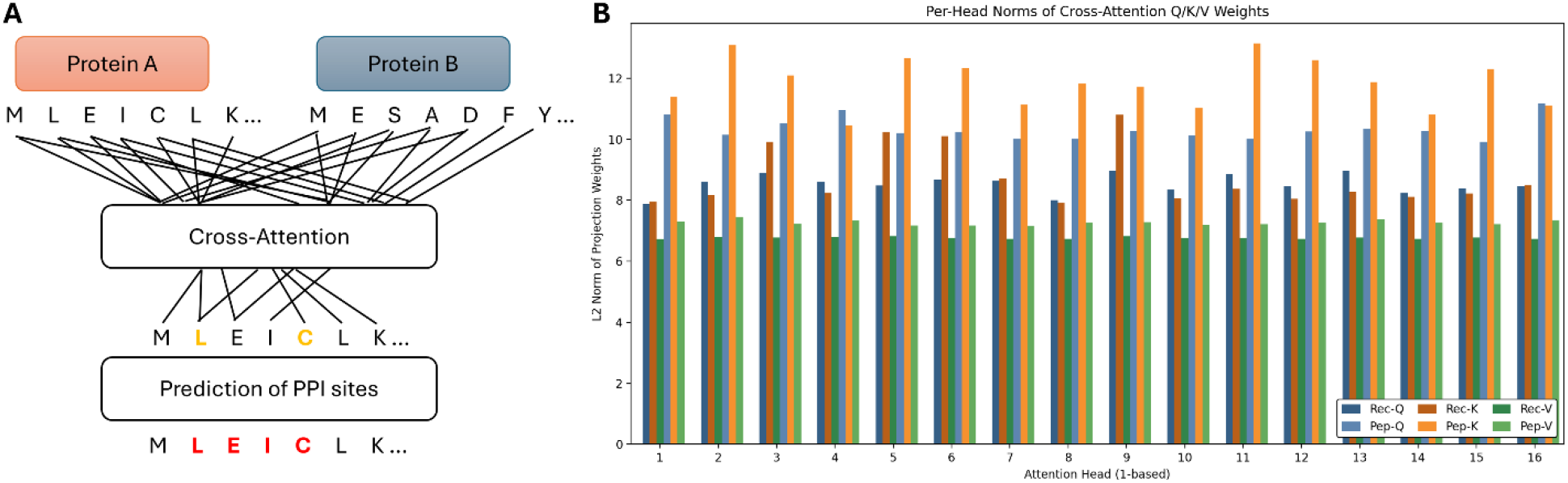
Cross-attention framework for partner-specific PPI site prediction and learned model statistics. **A.** Conceptual overview of the cross-attention mechanism. Residue-level embeddings of Protein A (query) and Protein B (partner) are provided as input. A cross-attention layer allows each residue in Protein A to attend to all residues in Protein B, generating partner-conditioned contextual representations. The resulting embeddings are processed by an MLP to yield residue-level interaction probabilities. **B**. Per-head L2 norms of Query (Q), Key (K), and Value (V) projection weights for receptor-centric (Rec) and peptide-centric (Pep) models. Both architectures share identical capacity (∼8.2 M parameters). Peptide-centric model displays systematically higher and more variable norms, particularly for Key projections, reflecting specialization for longer contiguous motifs typical of peptide interfaces, whereas the receptor-centric model shows lower and more uniform norms across heads, consistent with the dispersed and structurally heterogeneous nature of receptor interface patches.

Both receptor- and peptide-centric models share an identical architecture containing 8.20 million parameters, with approximately 80% of the total capacity allocated to the cross-attention block (Q/K/V projections, output projections, and LayerNorm) and the remaining ∼20% devoted to the classifier MLP (**Table 2**). This allocation underscores that the dominant representational burden lies in modeling partner-conditioned residue interactions rather than single-sequence features.

**Table 2.**
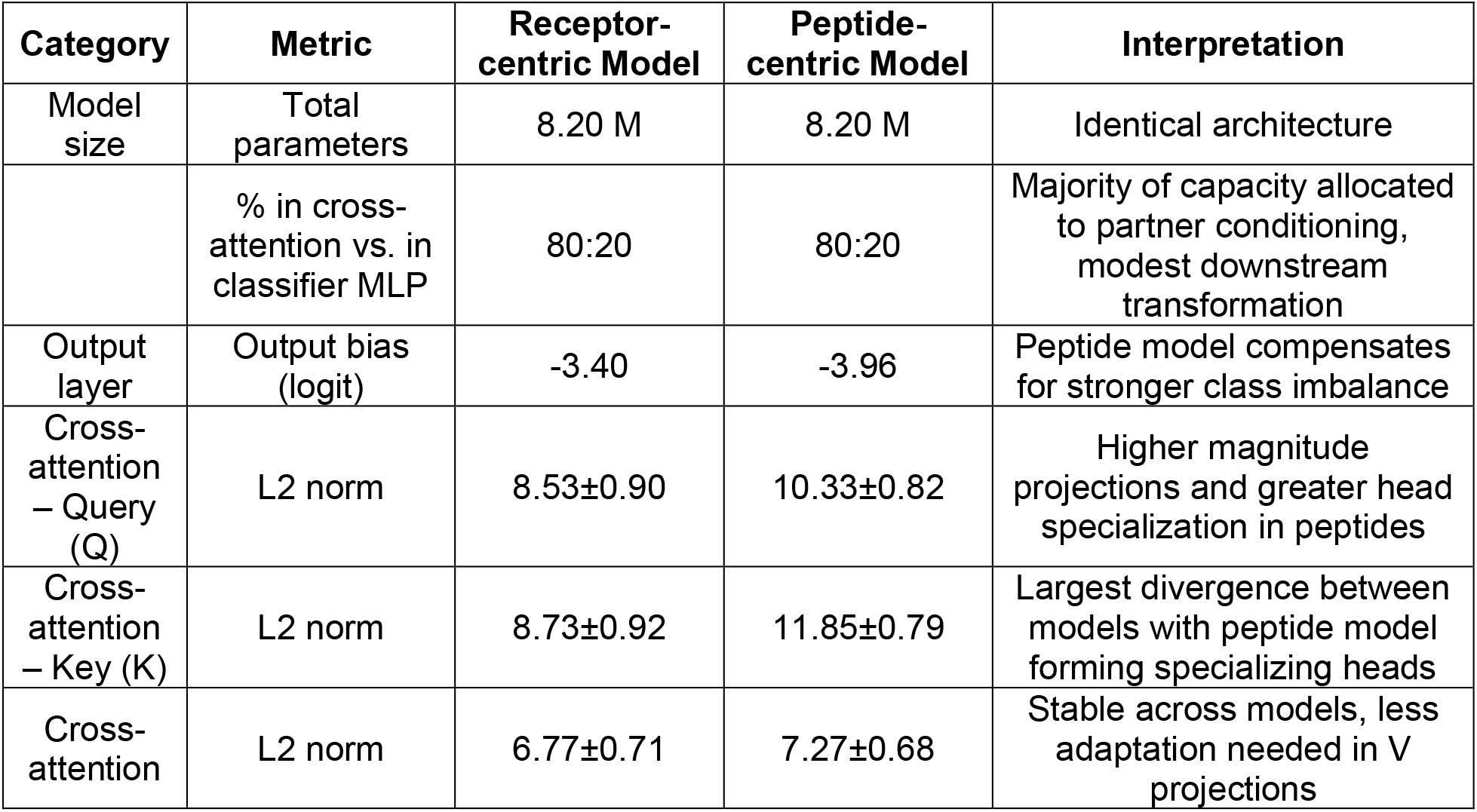
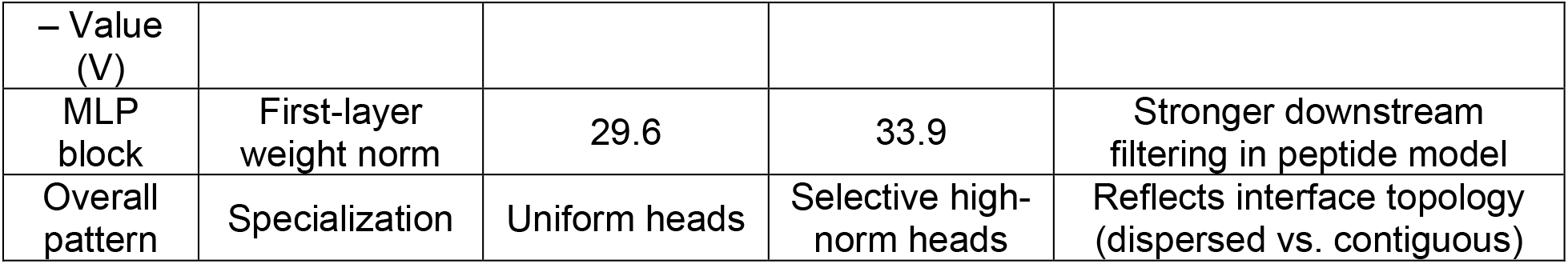
Summary of learned parameters and projection norms in receptor- and peptide-centric cross-attention models.

| Category | Metric | Receptor-centric Model | Peptide-centric Model | Interpretation |
| --- | --- | --- | --- | --- |
| Model size | Total parameters | 8.20 M | 8.20 M | Identical architecture |
|  | % in cross-attention vs. in classifier MLP | 80:20 | 80:20 | Majority of capacity allocated to partner conditioning, modest downstream transformation |
| Output layer | Output bias (logit) | -3.40 | -3.96 | Peptide model compensates for stronger class imbalance |
| Cross-attention – Query (Q) | L2 norm | 8.53±0.90 | 10.33±0.82 | Higher magnitude projections and greater head specialization in peptides |
| Cross-attention – Key (K) | L2 norm | 8.73±0.92 | 11.85±0.79 | Largest divergence between models with peptide model forming specializing heads |
| Cross-attention | L2 norm | 6.77±0.71 | 7.27±0.68 | Stable across models, less adaptation needed in V projections |
| – Value (V) |  |  |  |  |
| MLP block | First-layer weight norm | 29.6 | 33.9 | Stronger downstream filtering in peptide model |
| Overall pattern | Specialization | Uniform heads | Selective high-norm heads | Reflects interface topology (dispersed vs. contiguous) |

Quantitative inspection of trained checkpoints revealed consistent and biologically meaningful differences between the receptor- and peptide-centric models (**Table 2**). The peptide-centric model exhibited a more negative output bias (−3.96 vs. −3.40), consistent with the lower prevalence of positive interface residues in peptides and their higher class imbalance (**Table 1**). Analysis of per-head L2 norms showed systematically higher and more variable projection magnitudes in peptide-centric cross-attention, particularly for Key projections (mean 11.85 vs. 8.73 in receptors) and Query projections (mean 10.33 vs. 8.53). Value projections were more stable across models (means 7.27 vs. 6.77), indicating that most adaptive specialization occurs in how attention patterns are computed rather than how information is passed forward.

Consistent trends were observed in the first MLP layer, where peptide-centric weights exhibited larger norms (mean 33.9 vs. 29.6), reflecting stronger downstream transformations (**Table 2**). Together, these statistics support a mechanistic interpretation: the peptide-centric model develops a subset of high-magnitude, specialized attention heads that selectively amplify signals corresponding to longer contiguous motifs enriched in intrinsic disorder, features characteristic of short linear motif (SLiM)-mediated peptide interfaces. By contrast, the receptor-centric model maintains lower and more uniform magnitudes across heads, aligning with the more dispersed and patch-like interface organization found in structured receptor surfaces.

Details of the SPPIDER-seq implementation, including the web-based workflow, are provided in Supplementary Materials SD3.

### 3.2. Model Performance and Benchmarking against AlphaFold3

We evaluated both models on blind test subsets stratified by receptor- and peptide-centric contexts (RC-BS359 and PC-BS536, respectively). The receptor-centric model achieved an AUROC of 0.790 and a PRAUC of 0.212 (**Figure 2A–B**) whereas the peptide-centric model – AUROC of 0.797 and PRAUC of 0.205 (**Figure 2C–D**).

**Figure 2.**
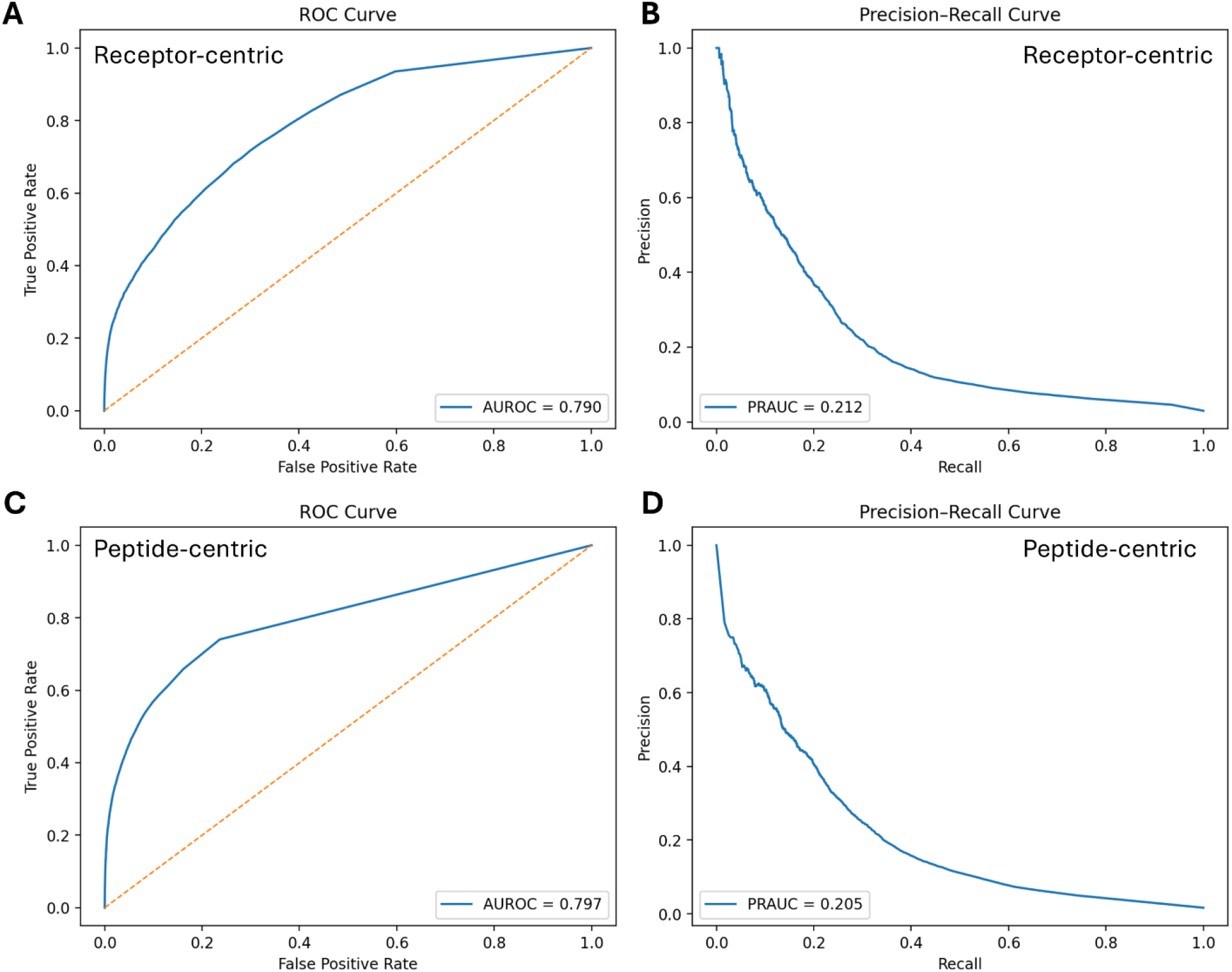
Performance of cross-attention models on blind test sets. **A–B.** ROC and precision– recall curves for the receptor-centric model based on the RC-BS359 blind set. **C–D**. ROC and precision–recall curves for the peptide-centric model based on the PC-BS536 blind set.

**Table 3** summarizes the detailed performance of the cross-attention models and AlphaFold3 (AF3) across corresponding subsets. Two primary trends emerged. First, the influence of the partner sequence varied between models. The peptide-centric model relied heavily on partner-specific information: replacing partner sequences with scrambled counterparts led to a pronounced performance drop (from MCC=0.269 down to 0.049; rows 7 vs. 10). Scrambled counterparts refer to partner sequences in which the amino acid order is randomly permuted while preserving residue composition, thereby disrupting all biologically meaningful motifs and interaction patterns. In contrast, the receptor-centric model was less dependent on partner identity (MCC=0.252 vs. 0.223; rows 1 vs. 4), suggesting that its predictions are driven primarily by structural cues intrinsic to the receptor embedding. This difference likely reflects the information content of interaction motifs: peptide SLiMs carry limited predictive signal on their own and require partner context, whereas receptor sites can be inferred from structural and surface exposure patterns.

**Table 3.**
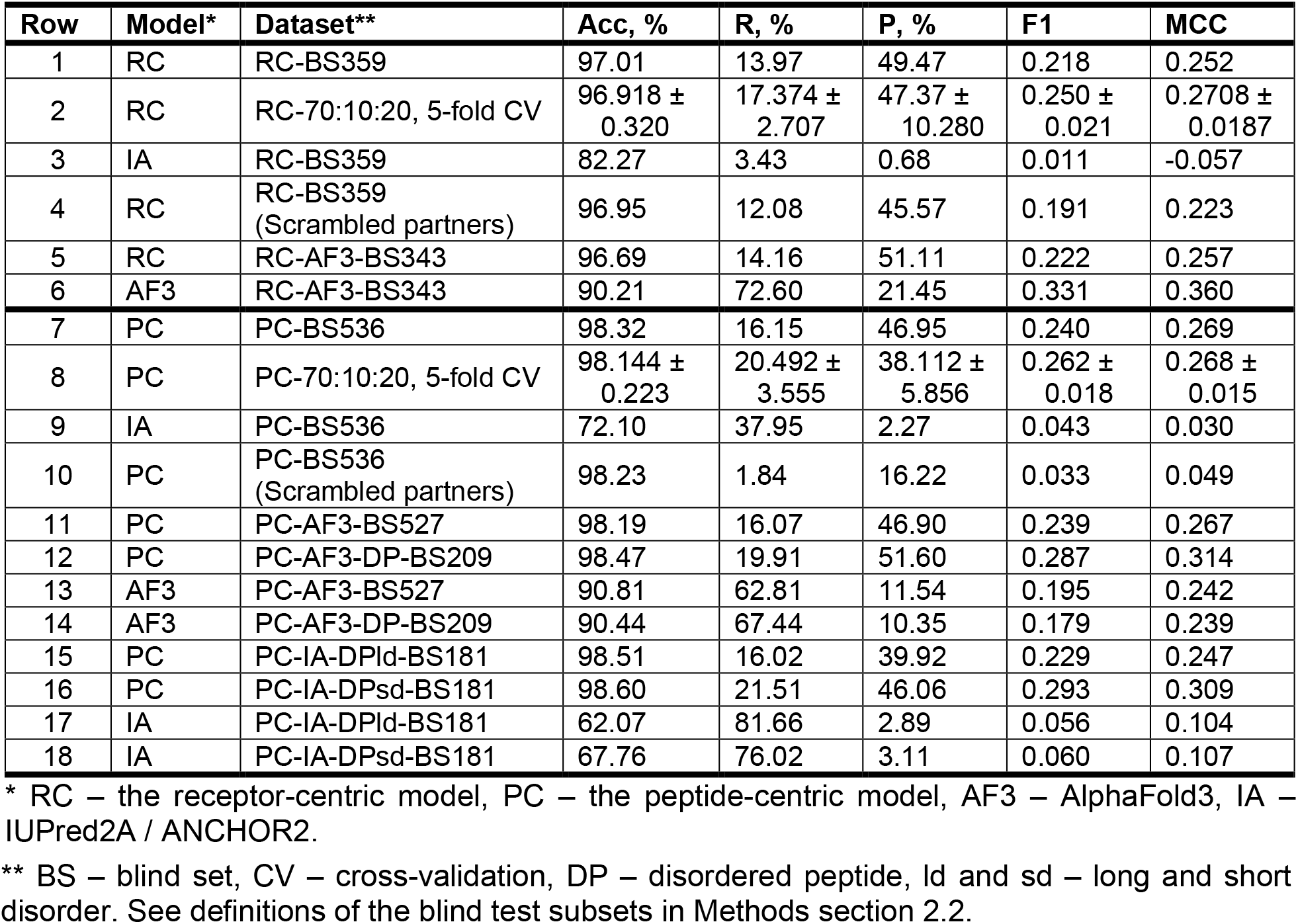
Performance of the cross-attention models and AlphaFold3 on the blind sets.

Second, AlphaFold3 tended to overpredict interfaces, identifying 60–70% of true positives but with precision of only 10–20%, indicating a high false-positive rate. While AF3 performed better than the receptor-centric model in structured datasets (MCC=0.360 vs. 0.257; rows 6 vs. 5), it struggled with small (MCC=0.242 vs 0.267; rows 13 vs. 11) and especially disordered peptide interfaces (MCC=0.239 vs. 0.314; rows 14 vs. 12). This limitation likely arises from AF3’s training bias toward well-structured complexes in the Protein Data Bank, which emphasizes stable domain–domain interactions and underrepresents transient, disordered contacts.

On the full receptor-centric blind set (RC-BS359; row 3), the IUPred2A/ANCHOR2 framework exhibited very low recall (3.43%) and precision (0.68%). On the peptide-centric blind set (PC-BS536; row 9), performance was characterized by substantially higher recall (37.95%) but similarly low precision (2.27%). Overall, the near-random performance observed on both datasets (MCC = −0.057 and 0.030, respectively) indicates that ANCHOR2 has limited ability to accurately localize interface residues when applied to full-length protein contexts.

When the peptide-centric blind set was restricted to proteins with PPI sites located within disordered regions, as defined by IUPred2A (PC-IA-DPld-BS181 and PC-IA-DPsd-BS181), ANCHOR2 performance improved in terms of sensitivity, achieving high recall but still very low precision, with MCC values only modestly above random (81.66%, 2.89%, 0.104 and 76.02%, 3.11%, 0.107 for the “long disorder” and “short disorder” definitions, respectively; rows 17 and 18). In contrast, SPPIDER-seq maintained substantially higher precision while preserving comparable recall and MCC to its performance on the full blind set (MCC = 0.247 and 0.309; rows 15 and 16). These results underscore the complementary characteristics of the two approaches: ANCHOR2 is highly sensitive to disordered binding regions but lacks specificity, whereas SPPIDER-seq achieves more precise and balanced residue-level predictions, particularly in biologically relevant settings where both ordered and disordered interfaces coexist. The SPPIDER-seq *vs*. ANCHOR2 contrast highlights that sequence-only disorder-based predictors capture general binding propensity but lack the contextual specificity required for precise interface localization.

### 3.3. Application of the Models to the PPI Site Prediction in TP53

TP53 encodes the transcription factor p53, a central regulator of cellular stress responses and the most frequently mutated gene in human cancer. Activated by diverse insults, p53 orchestrates transcriptional programs controlling cell-cycle arrest, DNA repair, apoptosis, and immune surveillance, thereby acting as a “guardian of the genome” (Hernández Borrero & El-Deiry, 2021; Kastenhuber & Lowe, 2017). Somatic TP53 mutations occur in ∼50% of tumors and include recurrent hotspot missense variants linked to characteristic mutational signatures, therapy resistance, and poor prognosis; germline mutations cause cancer-predisposition syndromes such as Li–Fraumeni (Muller & Vousden, 2013; Perri et al., 2016). Many mutant proteins also acquire gain-of-function activities that reprogram transcription, remodeling, and signaling networks (Muller & Vousden, 2013, 2014).

Structurally, p53 contains a folded DNA-binding core flanked by intrinsically disordered N- and C-terminal regions that are densely modified and highly interactive (Joerger & Fersht, 2010; Uversky, 2016; Xue et al., 2013). These IDRs harbor numerous short linear motifs that recruit E3 ligases (MDM2/MDM4), co-activators (p300/CBP), chromatin regulators, DNA-damage factors, and viral or oncogenic proteins, allowing p53 to integrate upstream cues and modulate target-gene selection (Joerger & Fersht, 2010; Uversky, 2016; Xue et al., 2013). Network-level analyses highlight p53 as a high-degree hub whose disease-associated mutations and regulatory sites frequently map to predicted disorder-based binding regions (Kastenhuber & Lowe, 2017; Uversky, 2016).

Accurately pinpointing p53 interaction sites is therefore critical for interpreting how mutations may rewire its interactome. Such knowledge also informs therapeutic strategies aimed at stabilizing wild-type p53 or disrupting oncogenic complexes (Hernández Borrero & El-Deiry, 2021; Joerger & Fersht, 2010; Muller & Vousden, 2014; Uversky, 2016). In this context, the 430 high-quality human TP53 interactions in the HINT database (Das & Yu, 2012), spanning 341 partners, provide a biologically rich testbed for partner-aware, residue-level PPI site prediction.

**Figure 3** reveals that predicted PPI hotspots are distributed nonuniformly along the p53 sequence, and these clusters align closely with its known domain architecture. According to NCBI CDD (Wang et al., 2023) annotation (**Figure 3A**), p53 comprises two N-terminal transactivation domains (TAD I: residues 6–30; TAD II: 35–59), a central DNA-binding domain (DBD: 99–289) containing a dimerization interface at 177–181, and a C-terminal oligomerization/tetramerization domain (319–358). The peptide-centric model predicts PPI site signals around residue positions ∼15–55, ∼330–350, and ∼380–393, whereas the receptor-centric model shows elevated median probabilities near ∼95–110, ∼120–130, ∼210–250, and ∼340–350 (**Figure 3B**). A substantial fraction of interaction partners exceed the ≥ 0.10 probability threshold in these same regions, suggesting recurrent engagement across diverse interaction contexts (**Figure 3C**). The cluster around the position 100 lies at the N-terminal boundary of the DBD and may correspond to interfaces with co-activators or regulatory proteins acting near the TAD/DBD junction. The strong signal between ∼220–250 aa falls within the DBD, implying that many partners engage surfaces on the folded core rather than only disordered flanking motifs. The pronounced cluster at ∼380– 393 aa spans the C-terminal regulatory region, consistent with frequent partner interactions that modulate oligomerization or bind post-translationally modified C-terminal tails, such as the cancer-testis antigen MORN3, reported to suppress p53 activity (Liang et al., 2018). Notably, the peptide-centric model produces stronger and broader predictions in the distal C-terminal segment, suggesting that many partner interactions are mediated by short linear motifs or disordered tails rather than by the well-folded receptor core. Together, these results indicate that the site-prediction models recover biologically plausible interface distributions that capture both structured-domain contacts and motif-mediated interactions in disordered regions, and that the human p53 interactome predominantly exploits two major interface zones: within the DBD and the C-terminal regulatory region, beyond the canonical N-terminal transactivation domains. Of note, PPI information from human p53 was effectively absent during both model training and validation. The only p53-containing complex included in the training subset of the receptor-centric model was PDB entry 7DVD, which captures the interaction between p53 (chain B, the “receptor”) and the pro-apoptotic Bcl-2–binding component 3, PUMA (chain E, the “ligand”). This interface involves only six residues within the p53 DNA-binding domain (S95, S96, V97, R209, N210, and F212), representing a small, localized surface patch on the folded core. No p53-related complexes appeared in any subset of the peptide-centric model. Consequently, the TP53 hotspot patterns identified by both architectures cannot be attributed to memorization of structural templates or inadvertent data leakage. Instead, they arise from generalizable interaction principles learned from diverse non-p53 protein pairs, reinforcing the biological plausibility and robustness of the predicted interface distributions. Collectively, the receptor-centric and peptide-centric models reveal largely distinct interaction profiles that overlap primarily within the TP53 tetramerization region, where different segments of the protein function alternately as receptor or ligand within the oligomeric assembly.

**Figure 3.**
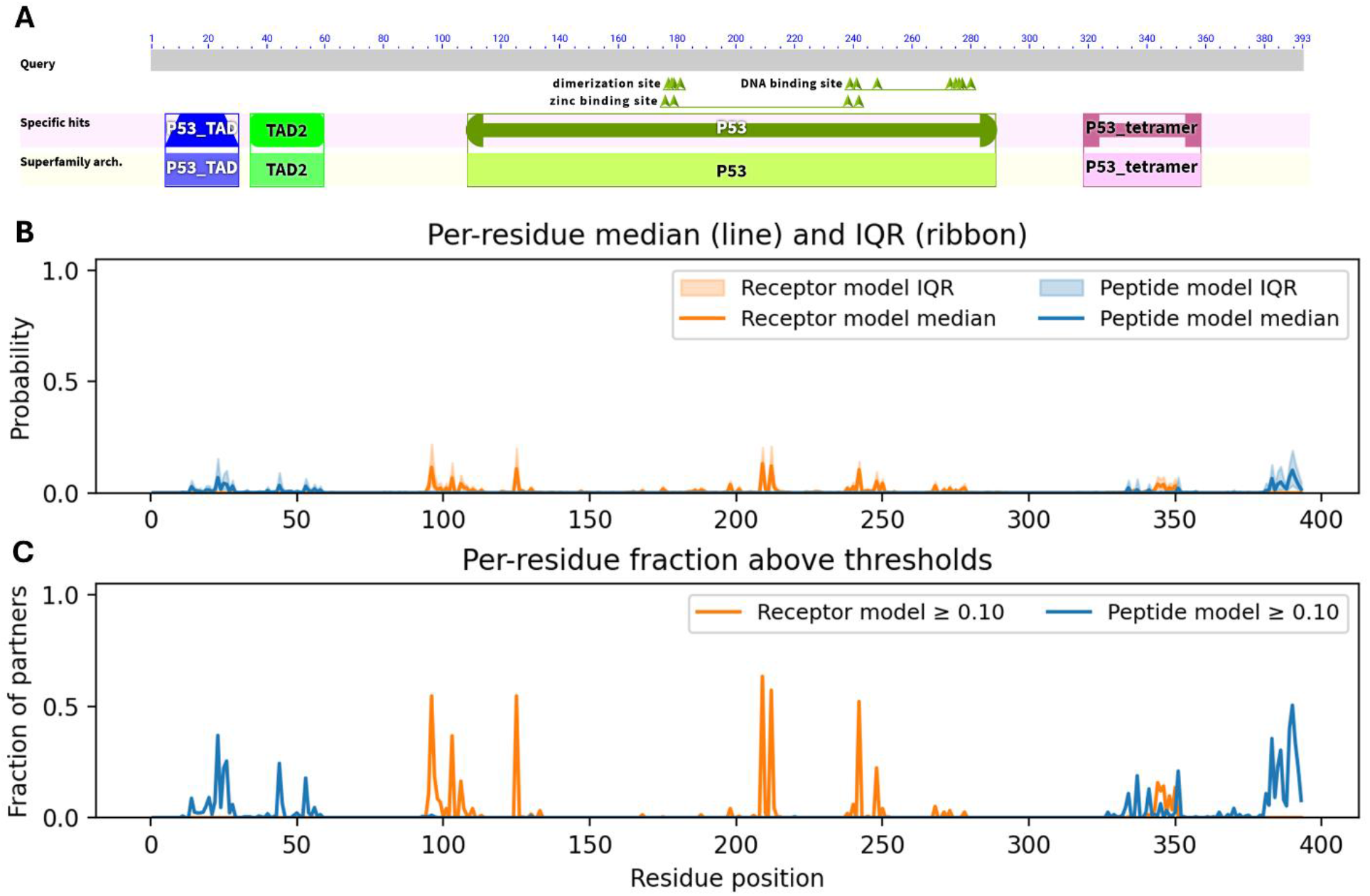
Partner-specific and model-specific patterns of predicted PPI site usage along the human TP53 sequence. **A.** NCBI CDD-based domain architecture and conserved functional regions of TP53, including transactivation domains (TAD1/2), DNA-binding and zinc-binding sites, and tetramerization motifs, shown to provide structural context for interpreting the prediction profiles. **B**. Per-residue distributions of predicted interaction probabilities across 341 experimentally validated TP53 partners, summarized as median scores (lines) and interquartile ranges (ribbons). These distributions illustrate both the magnitude and variability of partner-conditioned interface predictions. **C**. Fraction of partners for which each residue attains a predicted interaction probability ≥ 0.10, representing how frequently a given position is predicted to participate in binding across diverse interaction contexts.

## 4. Discussion

By explicitly modeling both interacting partners and allowing dynamic attention between them, the proposed cross-attention framework overcomes a major limitation of traditional single-input PPI site predictors. Previous models typically treated proteins in isolation, relying on static sequence or structural descriptors (Tang et al., 2023; Xue et al., 2015). Our architecture introduces partner-aware contextualization, enabling residue-level conditioning on the interacting partner and thereby improving localization of transient or disordered interfaces. This dynamic representation aligns with recent trends emphasizing contextualized embeddings and inter-protein co-representation learning (Liu et al., 2025; Si & Yan, 2024).

The enrichment of coil-based motifs in peptide-centric predictions reinforces the “disorder– function” paradigm, in which intrinsically disordered regions (IDRs) act as flexible scaffolds mediating transient binding through short linear motifs (SLiMs) (Ali et al., 2020; Seo & Kim, 2018; Wright & Dyson, 2015). Conversely, the receptor-centric model reproduces canonical features of structured-domain interactions, including α-helical and β-sheet surface patches typical of stable complexes (Chakrabarti & Bhattacharyya, 2007; Jones & Thornton, 1996). These complementary strengths indicate that the cross-attention mechanism learns biologically relevant structural priors from PLM embeddings without explicit structural supervision, a behavior consistent with emergent structure-awareness previously observed in ESM and ProtT5 embeddings (Elnaggar et al., 2022; Lin et al., 2023).

Although AlphaFold3 currently defines the state of the art for complex structure prediction (Abramson et al., 2024), our benchmarking reveals that it systematically overpredicts interface residues, particularly in disordered or peptide-mediated interactions. This likely reflects the evolutionary and structural bias of AF3 toward well-folded, globular complexes, inherited from the Protein Data Bank. In contrast, our partner-aware models remain robust when structural order is limited or transient – precisely where AF3 struggles. Similar behavior has been noted in comparative evaluations of deep-learning structure predictors versus disorder-centric binding models (Basu et al., 2023; Lee et al., 2024; Majila et al., 2026).

Inspection of learned attention weights provides interpretable clues about residue co-dependencies and partner-specific communication patterns. The peptide-centric model exhibits specialized attention heads with amplified Key/Query norms, resembling selective recognition modules tuned to SLiM-like motifs, whereas the receptor-centric model displays distributed attention indicative of broader interface patches. Such emergent specialization mirrors hierarchical binding principles observed in experimental PPI networks, where structured receptors recognize recurring motifs across multiple partners (Landgraf et al., 2004; Stein et al., 2009; Uversky, 2016).

The TP53 case study illustrates how partner-aware PPI prediction can map context-specific binding zones across multifunctional hub proteins. The recovered interface clusters coincide with functionally critical regions: transactivation motifs, DNA-binding core, and C-terminal regulatory tail, where cancer-associated mutations and post-translational modifications frequently occur (Kastenhuber & Lowe, 2017; Liang et al., 2018). Such residue-level maps can inform hypotheses regarding how specific mutations rewire the interactome or disrupt transcriptional regulation, complementing ongoing efforts in cancer variant interpretation (Joerger & Fersht, 2010).

Despite these strengths, several limitations of the current models warrant consideration. First, because experimentally resolved PPI interfaces that depend explicitly on post-translational modifications (PTMs) are sparse in public datasets, the models are effectively trained on PTM-agnostic interaction data. As a result, SPPIDER-seq cannot reliably distinguish PTM-dependent interfaces, such as phospho-switches or acetylation-regulated binding sites, even though protein language model embeddings may implicitly capture sequence features associated with PTM propensity. Explicit integration of curated PTM–PPI datasets will be required to model such regulatory interactions. Second, although the method accurately localizes interaction-prone regions, the absolute predicted probabilities remain largely low. These conservative probability distributions lead to reduced recall at conventional thresholds and may underrepresent weak, transient, or context-specific binding. Third, because training labels are derived from static receptor–peptide complexes, the models do not fully capture stoichiometric diversity or cooperative binding, such as multi-interface assemblies, allosterically coupled contacts, or competitive binding between partners. Future work incorporating PTM-aware supervision, improved calibration of predicted probabilities, and training on broader classes of interaction data may help address these constraints.

Together, these findings demonstrate that the cross-attention paradigm bridges a critical gap between sequence-based and structure-based models, yielding biologically interpretable, scalable, and partner-aware predictions of protein–protein interaction sites. Importantly, partner-aware prediction shifts the problem from identifying all possible interfaces to prioritizing context-specific interaction hypotheses, which is more aligned with experimental validation workflows.

By combining ESM-2 embeddings with cross-attention, SPPIDER-seq establishes a flexible foundation for proteome-scale, context-aware PPI site prediction. Potential extensions include: (1) Variant impact modeling – assessing mutation-induced changes in partner-conditioned interaction probability; (2) Network reconstruction – integrating residue-level scores to refine interactome maps or predict rewired signaling in disease; (3) Drug discovery – identifying transient or cryptic hotspots suitable for small-molecule or peptide inhibitors; and (4) Methodological advances – extending to multi-partner attention, integration with structural priors (AlphaFold3 geometries), or fine-tuning with experimental datasets such as deep mutational scanning.

## Acknowledgements

This work was supported by the National Institutes of Health [1R01HL176858 to A.P., 1R35HL171346 to J.C.].

## Notes

https://github.com/aporollo-lab/SPPIDER-seq

https://doi.org/10.5281/zenodo.19835990

### Competing Interest Statement

The authors have declared no competing interest.

### Summary of Updates

This revised version substantially expands and clarifies the SPPIDER-seq framework for partner-aware protein-protein interaction site prediction. Major updates include implementation of 5-fold grouped cross-validation with non-overlapping UniProt-based splits to improve robustness assessment and reduce partition-specific bias. The dataset construction and redundancy filtering procedures were clarified, including explicit explanation that full-length UniProt parent proteins rather than isolated peptide fragments were used during training and benchmarking. The revised manuscript now includes comprehensive comparison against the sequence-based disorder and binding-region predictors IUPred2A and ANCHOR2.

