## Supplementary Materials for "SPPIDER-seq: Sequence-based partner-aware predictor of protein-protein interaction sites"

#### Supplementary Data SD1. Analysis of Protein-Protein Interaction Interfaces

Analysis of interface residues in the overall sets of receptor–peptide complexes revealed distinct organizational and structural characteristics. At a qualitative level, receptor-centric interfaces tended to form multiple dispersed patches, whereas peptide-centric interfaces typically comprised broader and more contiguous stretches (**Figure S1A**). Quantitative assessment of motif length distributions confirmed this observation (**Figure S1B**). Receptor interfaces were dominated by short, discontinuous motifs, while peptide ligands exhibited longer continuous segments of binding residues. Although peptide interfaces are smaller in overall size (**Figure S1B**), their extended contiguous binding stretches span more residues on average, consistent with the presence of short linear motif (SLiM)-like elements embedded in disordered regions.

To further examine the structural context of these sites, we analyzed single-chain 3D models from the AlphaFold2 database (**Figure S1C**). In receptor proteins, interface residues were distributed across multiple secondary-structure elements, with α-helices and β-strands accounting for approximately 45% and 20% of the sites, respectively. In contrast, peptide interfaces were strongly enriched in coil conformations (~61%). Collectively, these results underscore fundamental organizational differences between receptor and peptide interfaces and highlight the prominent role of SLiMs in mediating transient and regulatory protein interactions.

These observations motivated the development of two complementary, partner-aware predictors: a receptor-centric model, optimized for structured protein surfaces, and a peptide-centric model, tailored for interactions mediated by extended, unstructured motifs.

**A**

```
Rec1: 00011010101100110010000000000000000001001101100100000
Rec2: 00000000000000001001100110110010000000000000000000000
Rec3: 00110100000000000000000011111111000000000001001010100
Rec4: 01001000100000000000000000100100011000000000000000000

Lig1: 00000000000111111111111110000000000000000000000000000
Lig2: 00000000000000000000001111111111111000000000000000000
Lig3: 00000000000000000000000000000111111011111111111110000
Lig4: 00000001111000000000000100011110000000000000000000000
```

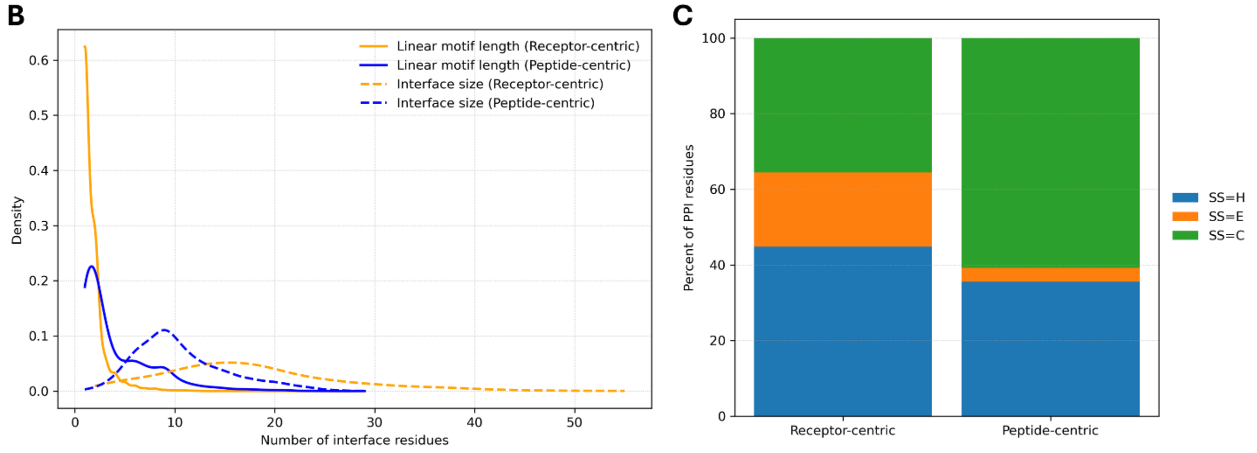


**Figure S1. Distinct structural and organizational properties of interface residues in receptor–peptide complexes. A.** Representative binary sequence profiles illustrating interface residues in receptor proteins (Rec) and peptide ligands (Lig) curated from the BioLiP database. **B.** Density distributions of interface residue counts, including both the lengths of continuous interface motifs (solid lines) and the total interface sizes (dashed lines), shown for the receptor-centric (RC-Overall) and peptide-centric (PC-Overall) datasets (Section 2.2 of the main manuscript). The comparison reveals enrichment of short, contiguous motifs in peptides, whereas receptors exhibit larger, more distributed interfaces. **C.** Secondary-structure composition of interface residues derived from full-length AlphaFold2 models, showing a higher prevalence of α-helices and β-strands in receptors and coil conformations in peptides. SS: H – α-helix, E – β-strand, C – coil.

**Supplementary Data SD2. Evaluation Metrics**

The following threshold-dependent metrics were used:

- Accuracy, *Acc* – the proportion of correctly classified residues:

$$Acc = \frac{TP + TN}{TP + TN + FP + FN}$$

- Recall (Sensitivity), *R* – the proportion of true interface residues correctly identified:

$$R = \frac{TP}{TP + FN}$$

- Precision (Positive Predictive Value), *P* – the proportion of predicted interface residues that are true:

$$P = \frac{TP}{TP + FP}$$

- F1-score, *F1* – the harmonic mean of precision and recall:

$$F1\text{-}score = \frac{2 \cdot \mathrm{Precision} \cdot \mathrm{Recall}}{\mathrm{Precision} + \mathrm{Recall}}$$

- Matthews Correlation Coefficient, *MCC* – a balanced measure of classification quality, robust to class imbalance:

$$MCC = \frac{TP \cdot TN - FP \cdot FN}{\sqrt{(TP + FP)(TP + FN)(TN + FP)(TN + FN)}}$$

Here, TP (true positives), TN (true negatives), FP (false positives), and FN (false negatives) represent residue-level outcomes of interface classification.

In addition, two threshold-independent metrics were included. First is Area Under the Receiver Operating Characteristic Curve (AUROC) – quantifies the model's ability to discriminate between interface and non-interface residues across all thresholds. A value of 1.0 indicates perfect separation, while 0.5 corresponds to random guessing. Second is Area Under the Precision–Recall Curve (PRAUC) – reflects the trade-off between precision and recall across thresholds. PRAUC is particularly informative for imbalanced PPI datasets dominated by negatives, emphasizing model performance on the positive class (interface residues).

All metrics were computed exclusively on the query (focal) proteins – the proteins for which interface residues were predicted. The partner proteins, although used to provide contextual embeddings via cross-attention, were not evaluated directly. Of note, the AUROC and PRAUC metrics could not be used to benchmark AlphaFold3, since AF3 does not output residue-level probabilities. Instead, the interface annotations were derived from the 3D structures of protein complexes predicted in the top-ranked AF3 models, and binary residue states (interface *vs*. non-interface) were used for evaluation.

### Supplementary Data SD3. Software Architecture and Associated Utilities of SPPIDER-seq

SPPIDER-seq is implemented in PyTorch and distributed as a Jupyter Notebook designed for deployment on the Google Colab platform, with support for both GPU- and CPU-based execution (**Figure S2**). The notebook provides an interactive interface in which users may submit one or multiple query and partner protein sequences. When multiple sequences are supplied, the framework automatically performs predictions for all query–partner pairwise combinations.

Residue-level predictions are reported in two complementary formats. First, graphical outputs display predicted interaction probabilities along the query sequence, with results from the receptor-centric and peptide-centric models overlaid to facilitate visual inspection of putative PPI site regions. Second, tab-delimited text files provide numerical probability values for each residue, enabling straightforward downstream processing and integration with external analytical pipelines.

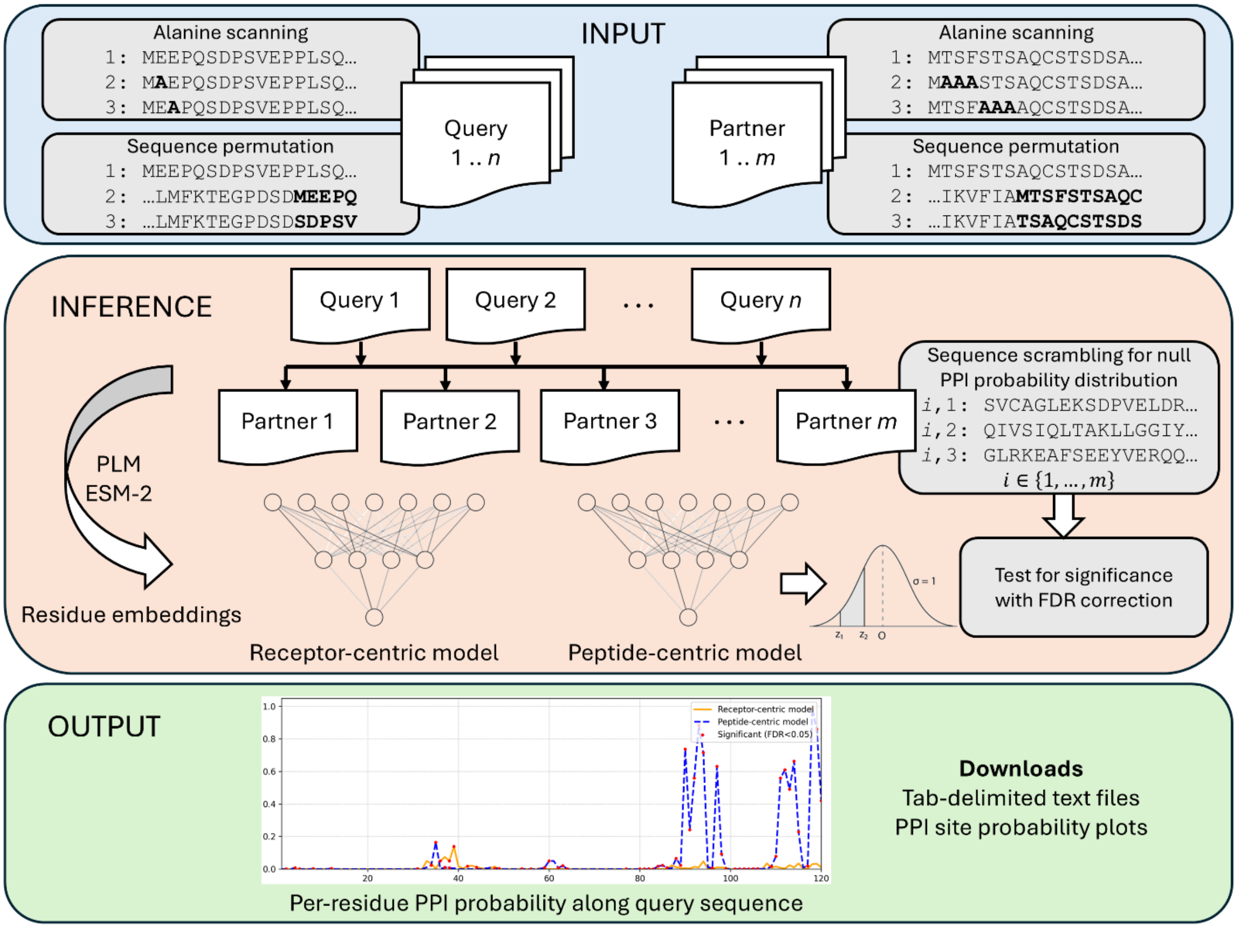


**Figure S2.** Web-based workflow and analytical utilities of SPPIDER-seq. Schematic overview of the SPPIDER-seq pipeline illustrating data flow from input to output. **Input** (top): one or multiple query proteins (1 … *n*) and interaction partners (1 … *m*) are provided by the user. Optional experimental-design utilities (grey background) allow generation of perturbed sequence sets, including alanine-scanning variants and sequence permutation, for both queries and partners. **Inference** (middle): query–partner all-by-all combinations are evaluated. Protein sequences are first converted into context-aware, per-residue embeddings using a frozen protein language model (ESM-2). These embeddings are processed by two complementary cross-attention models, a receptor-centric and a peptide-centric model, to produce partner-conditioned residue-level interaction logits. An optional null-background analysis (grey background) can be invoked, in which partner sequences are scrambled to estimate a null distribution of PPI probabilities; observed residue scores are then assessed for statistical significance using z-score–based testing followed by Benjamini–Hochberg false discovery rate (FDR) correction. **Output** (bottom): results are reported as per-residue PPI probability profiles along the query sequence, with predictions from both models overlaid and statistically significant positions highlighted when applicable, and as downloadable tab-delimited text files containing numerical probability and significance values. Grey-shaded blocks denote optional, computationally intensive steps that are typically used for targeted analyses or experimental interpretation rather than routine large-scale inference.

An optional null-background analysis is implemented to assess the statistical significance of predicted PPI sites. When enabled, the notebook generates several predictions for each query in the context of multiple scrambled versions of the partner sequence. For each residue, the observed prediction is evaluated against the null distribution using a z-score–based test, followed by Benjamini–Hochberg correction to control the false discovery rate. Statistically significant positions are annotated both in the graphical outputs and in the accompanying text files. Because this procedure substantially increases computational cost, it is intended primarily for single query–partner analyses in Google Colab or for large-scale applications executed on local workstations or high-performance computing environments.

In addition to the core prediction workflow, the repository includes a collection of utility notebooks supporting experimental design and model interpretation, including partner sequence scrambling or permutation, and alanine-scanning–style *in silico* mutagenesis. These tools enable systematic interrogation of model sensitivity to partner identity and local sequence variation, thereby extending SPPIDER-seq from a predictive framework to an interpretable platform for generating mechanistic hypotheses about protein–protein interaction determinants.
